# A Test for Confounding in Coupling of Multimodal Neuroimaging Data

**DOI:** 10.64898/2026.09.14.750774

**Authors:** Yiyan Hao, Simon N. Vandekar, Aaron F. Alexander-Bloch, Theodore D. Satterthwaite, Brian R. White, Russell T. Shinohara

**Author notes:** These authors contributed equally as senior authors.

## Abstract

Multimodal neuroimaging studies often involve comparisons between brain maps. Recently, statistical methods have been proposed to quantify and assess spatial correspondence between two modalities. The simple permutation-based inter-modal correspondence (SPICE) test evaluates whether within-subject correspondence exceeds chance, where the null distribution is constructed by permuting subject labels for one modality. Despite its easy implementation and minimal spatial assumptions, the critical assumption underlying permutation analysis, that subjects are exchangeable under the null, may be violated when covariates such as age, sex, or disease status systematically alter brain map distributions. This violation can produce results analogous to Simpson’s paradox, where apparent population-level correspondence reflects between-group differences rather than genuine within-subject coupling. We propose a formal U-statistic-based test for such covariate effects, enabling both diagnostic evaluation of assumption violations and scientific discovery. Using synthetic and semi-synthetic neuroimaging data, we demonstrate well-controlled Type I error and high statistical power. We apply our method to test for confounding effects due to age and sex using real data from two pairs of imaging modalities in the Philadelphia Neurodevelopmental Cohort. Our framework increases the rigor and interpretability of intermodal coupling analyses, with broad implications for neuroimaging studies in heterogeneous populations, especially in developmental, aging, and disease-focused research.

## 1 Introduction

Advances in neuroimaging and large-scale initiatives have made subject-level brain maps across multiple modalities widely available, enabling researchers to link imaging phenotypes with cognition, aging, behavior, and disease (Jirsaraie et al., 2023; Kalpouzos and Persson, 2025; Elliott et al., 2018; Smith et al., 2021; Bethlehem et al., 2022; Cole et al., 2017; Kaufmann et al., 2019; Drysdale et al., 2017). A central question raised by these multimodal datasets is the spatial correspondence problem – assessing whether two brain maps show meaningful convergence, overlap, or topographic similarity. For example, correspondence between task activation maps, resting-state functional networks, and other functional annotations may reveal shared neural substrates of cognition (Smith et al., 2009; Mennes et al., 2010; Cole et al., 2014; Tavor et al., 2016; Cole et al., 2016); whereas correspondence between morphological features such as cortical thickness and surface area may provide insight into shared or distinct developmental, cellular, or genetic influences (Grasby et al., 2020; Seidlitz et al., 2018; van der Meer et al., 2023; Panizzon et al., 2009). A traditional approach is to visually compare the pair of maps side by side, which is intuitive but yields only qualitative conclusions. On the other hand, classical hypothesis testing procedures can provide quantitative inference, but they often rely on strong assumptions that are inappropriate for spatially structured brain data. For example, when testing with a parametric null such as the Student’s *t* distribution, the assumptions that model errors are independent and identically distributed across locations are violated due to the existence of spatial autocorrelation in neuroimaging data. Similarly, using non-parametric methods, one may obtain *p*-values by freely permuting map values across spatial locations, but the assumption that locations are exchangeable is also violated by the strong spatial autocorrelation. As a result, the null distribution may be composed of biologically implausible surrogate maps, leading to spuriously small *p*-values (Markello and Misic, 2021). These limitations have motivated the development of a growing class of spatially constrained statistical approaches.

The spin test (Alexander-Bloch et al., 2018) is a widely adopted approach that generates null models by applying random rotations to spherical projections of the cortical surface, a permutation framework that preserves spatial autocorrelation. Another class of methods, represented by BrainSMASH (Burt et al., 2020), instead simulates surrogate maps with matched spatial autocorrelation structure using variogram-based or parametric generative models. These spatial null approaches have been widely applied to test correspondence between structural, functional, transcriptomic, and neurotransmitter receptor maps (Hansen et al., 2022; Burt et al., 2018; Bazinet et al., 2023; Sun et al., 2025; Sydnor et al., 2025; Wu et al., 2025; Paquola et al., 2025; Liao et al., 2023), and their statistical properties and performance have been further benchmarked (Markello and Misic, 2021; Váša and Mišić, 2022). While effective for group-averaged or single-subject maps, most of these approaches do not naturally extend to subject-level paired data.

The simple permutation-based inter-modal correspondence (SPICE) test (Weinstein et al., 2021), on the other hand, assesses spatial correspondence by testing whether two neuroimaging modalities show stronger correspondence within the same individual than would be expected by chance. Unlike the aforementioned methods that operate on group-averaged maps and construct spatial null models, the SPICE test uses subject-level paired imaging data and generates a null distribution by permuting the subject labels for one of the two modalities, or equivalently shuffling subject pairings. The observed average within-subject correspondence statistic is then compared with its permuted counterparts to obtain a permutation-based *p*-value. Thus, the SPICE test is intuitive, easy to implement, and avoids strong assumptions about the brain’s spatial covariance structure.

A key assumption of the SPICE permutation framework is that subjects are exchangeable under the null hypothesis. However, brain maps are known to vary systematically with development, aging, sex, and disease status (Bethlehem et al., 2022; Lenroot and Giedd, 2012; Kaufmann et al., 2019). In such settings, individuals from different subpopulations may exhibit distinct spatial patterns within one or both modalities, such that unrestricted permutation across subjects may exchange maps whose distributions differ under the null. Consequently, the aggregate association differs from, or even masks or exaggerates, the associations present within relevant subgroups, resulting in a potential failure mode analogous to Simpson’s paradox (Simpson, 1951; Blyth, 1972). In the context of the SPICE test, the overall degree of intermodal correspondence may therefore reflect differences in subgroup composition, such as age-related or disease-related variation, rather than homogeneous subject-level coupling. To mitigate this ambiguity, we propose a new method to identify such covariate effects in the SPICE test.

This test serves two complementary purposes. First, as a diagnostic tool, it distinguishes genuine within-subject coupling from apparent coupling that arises solely due to difference between subgroups, and guides analysts on when the SPICE test should be applied conditionally instead of on the full sample. Second, it can be employed to identify covariates that modulate intermodal coupling, providing a principled way to characterize how biological or demographic factors shape the relationship between different brain features.

The rest of this article is organized as follows. Section 2 introduces basic notation for the SPICE test and defines the hypotheses of interest. Section 3 provides concrete examples using neuroimaging data from the Philadelphia Neurodevelopmental Cohort (PNC), which was used in the original formulation of SPICE, to illustrate the existence and consequences of confounding effects. Section 4 proposes a test for such covariate effects based on U-statistic theory. Section 5 presents simulation studies to evaluate the proposed methods using fully synthetic data and pseudo brain maps derived from real neuroimaging data. Section 6 presents an application of our proposed testing framework to examine intermodal coupling in the PNC dataset while accounting for age and sex effects, respectively. Section 7 gives concluding remarks.

## 2 The SPICE Test In The Presence Of Covariate Effects

### 2.1 Notation

Let *X_i_, Y_i_* ∈ ℝ*^p^* denote vectors representing spatial maps (e.g., neuroimaging data) in a common template space derived from the two modalities under comparison, observed for individuals *i* = 1*, …, n*. Suppose that there exists a binary covariate *G_i_*that categorizes each individual into one of two underlying sub-populations *A, B* with sample sizes *n_A_, n_B_*, respectively, where *n_A_* + *n_B_* = *n*. That is,

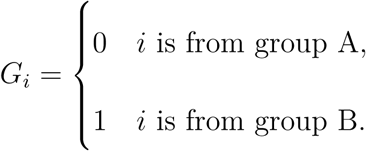

Let *ψ*(·, ·) be a symmetric measure of the intermodal correspondence between two vector maps. Without loss of generality, we use the Pearson correlation *ρ*(·, ·) as the correspondence measure for the following derivation.

### 2.2 Global SPICE test

Here, we restate the formulation of the SPICE test (Weinstein et al., 2021). As this method uses all subjects’ data without respect to other covariates, we refer to it as the global SPICE test. The null hypothesis states that the between-subject intermodal correspondence has the same distribution as the within-subject intermodal correspondence. We refer to this null hypothesis as the global null:

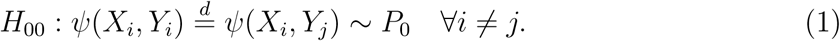

*H*_00_ is tested by comparing the observed test statistic

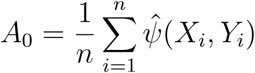

with

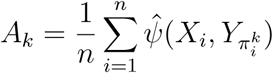

from permutations *k* = 1*, …, K*, where 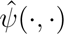 is the observed intermodal correspondence\ (e.g., sample correlation) and 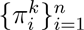 is a permutation of {*i* : *i* = 1*, …, n*}. Here, *A*_0_ gives the empirical mean of the left hand side of (1) using the observed subject-level data. *A_k_* estimates the mean of the right hand side of (1) under *H*_00_ through permutations. These lead to the SPICE *p*-value

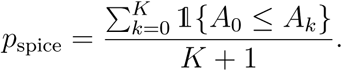

### 2.3 Conditional SPICE test

The global null hypothesis *H*_00_ is a statement about the marginal distributions, which does not take into account any potential dependence on the covariates. However, in many applications, one may be aware or suspicious of potential confounders whose value may affect the degree of intermodal coupling. For instance, morphometric features of the brain may develop with age or change over disease progression. Such a setting could result in estimates of coupling whose magnitude is attributable to these confounders as opposed to underlying intermodal relationships within participants. Rather, of greater interest in such a scenario are the conditional null hypotheses within each group as follows:

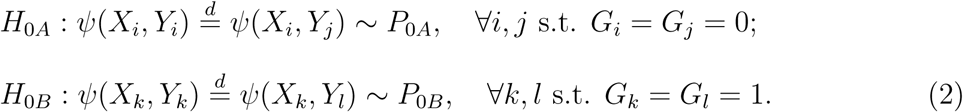

The permutation framework used by the SPICE test assumes exchangeability of data under the null *H*_00_: that swapping subject labels for maps of the second modality does not affect the joint distribution of (*X_i_, Y_i_*)*, i* = 1, · · · *, n*. This assumption is violated in the presence of a subject-level covariate that strongly influences the data distribution. In such cases, the corresponding conditional null hypotheses *H*_0_*_A_, H*_0_*_B_* can be tested by the SPICE test statistics computed with data from only the group being tested, with

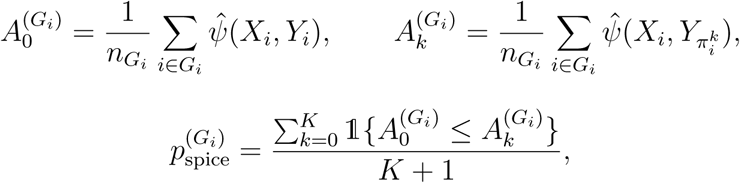

where 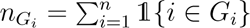. We refer to this procedure as the conditional SPICE test.

## 3 Motivating Study

### 3.1 Introduction

Before addressing how to know when it may be appropriate to use the conditional SPICE as compared to the current global SPICE, we first motivate the statistical problems that can arise from covariate effects in the context of the SPICE test. We used the same dataset and comparisons from the original formulation of the SPICE test by Weinstein et al. (2021). Neuroimaging data were taken from the Philadelphia Neurodevelopmental Cohort (PNC) (Satterthwaite et al., 2014, 2016), and we considered the same two sets of intermodal comparisons: cortical thickness against sulcal depth; cortical thickness against the 2-back *>* 0-back contrast from a fractal n-back working memory task (Frac2back). In addition to reproducing findings in Weinstein et al. (2021), we provide new insights into how coupling between imaging modalities may be confounded by age.

### 3.2 Data description

Collected through the University of Pennsylvania and Children’s Hospital of Philadelphia, the PNC provides a rich repository of subject-level multimodal MRI data in addition to cognitive, clinical, and genomic phenotypes from a large youth sample (Satterthwaite et al., 2014, 2016). It has been widely used to capture coordinated neurodevelopmental change across brain structure, functional connectivity, and structure–function coupling. Prior PNC-based studies have shown that deviations from normative maturational patterns are associated with cognition, executive function, attention, and psychiatric vulnerability (Satterthwaite et al., 2013; Erus et al., 2015; Kessler et al., 2016; Baum et al., 2017; Xia et al., 2018; Baum et al., 2020; Gur et al., 2025; Sydnor et al., 2025; Li et al., 2026; Luo et al., 2026).

Cortical thickness refers to the distance between the gray-white matter boundary and the pial surface, while sulcal depth quantifies the depth of cortical folds relative to the outer cortical surface. In MRI analysis, both measures are typically derived from structural T1-weighted images. Cortical thickness and sulcal depth capture complementary but coupled aspects of cortical morphology. Studies have shown that the cortex is generally thinner in sulcal fundi than on gyral crowns, suggesting that cortical thickness varies systematically with local folding topology (Vandekar et al., 2015). Moreover, this relationship is developmentally dynamic, making this pair of modalities a natural candidate to which our test for covariate (age) effect can be applied. Previous analyses of the PNC dataset found that cortical thinning is spatially patterned by sulcal-gyral location and that subject-level thickness-depth coupling shows heterogeneous, nonlinear age-related trajectories (Vandekar et al., 2015, 2016). Longitudinal adolescent data further suggest that cortical maturation includes coordinated flattening of the cortical surface, with sulci becoming shallower and wider as cortical thickness decreases (Alemán-Gómez et al., 2013).

The fractal n-back task, on the other hand, provides a functional measure of working memory. In this working-memory functional MRI (fMRI) paradigm, participants view abstract fractal images and respond when the current image matches one shown (n) trials earlier. Satterthwaite et al. (2013) found that working memory performance improved with development and was strongly associated with executive-network activation and default-mode deactivation, suggesting that n-back performance reflects functional maturation of executive systems. We do not expect a strong correspondence between structural morphology maps and task-activated cognitive maps.

We included all subjects from PNC that had all three modalities available and which passed quality requirements (*N* = 1347) (Shafiei et al., 2025). We analyzed data in fsaverage space (Fischl et al., 1999) with *p* = 163842 vertices.

### 3.3 Conditional SPICE tests reveal age-specific coupling

Following the analysis approach in Weinstein et al. (2021), we first performed the SPICE test using the entire sample. As in Weinstein et al. (2021), the global results suggested significant coupling between cortical thickness and sulcal depth but not between cortical thickness and Frac2back (Table 1).

**Table 1:** Results from global and conditional SPICE performed on a subset of the PNC dataset. For all comparisons, we used measures from the left hemisphere for each subject, and reported the observed correlations 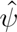 and corresponding permutation-based *p*-values. CT: cortical thickness; SD: sulcal depth.

| Age | N | CT vs SD |  | CT vs Frac2back |  |
| --- | --- | --- | --- | --- | --- |
| | | $p$ | $\hat{\psi}$ | $p$ | $\hat{\psi}$ |
| All | 1347 | < 0.001 | −0.147 | 0.950 | −0.051 |
| [8, 10) | 110 | < 0.001 | −0.108 | 0.221 | −0.032 |
| [10, 12) | 175 | < 0.001 | −0.131 | 0.720 | −0.048 |
| [12, 14) | 198 | < 0.001 | −0.142 | 0.990 | −0.041 |
| [14, 16) | 237 | < 0.001 | −0.149 | 0.740 | −0.041 |
| [16, 18) | 254 | < 0.001 | −0.153 | 0.610 | −0.066 |
| [18, 20) | 242 | < 0.001 | −0.163 | 0.500 | −0.060 |
| [20, 22) | 119 | < 0.001 | −0.163 | 0.520 | −0.056 |
| [22, 24) | 12 | < 0.001 | −0.164 | 0.061 | −0.078 |

We then divided the sample into 8 groups of consecutive age with increments of 2 years ([8, 10), [10, 12)*, …,* [22, 24)) to test for intermodal correspondence in each group separately. When examining the age groups separately, we identified interesting patterns in which both the observed test statistic and its null distribution shifted with age (Fig 1). The magnitude of the observed correlation between cortical thickness and sulcal depth increased monotonically as age increased. Their corresponding null distributions, constructed by permutations in the conditional SPICE tests, also moved in the same direction (Fig 1A). This co-movement of the observed statistic and its null indicates that age is a confounding covariate that changes the marginal distributions of cortical thickness and sulcal depth maps, which in turn shifts the range of correlation values attainable by chance alone. On the other hand, the correlation between cortical thickness and Frac2back reflected a non-monotone developmental effect (Fig 1B).

**Figure 1:**
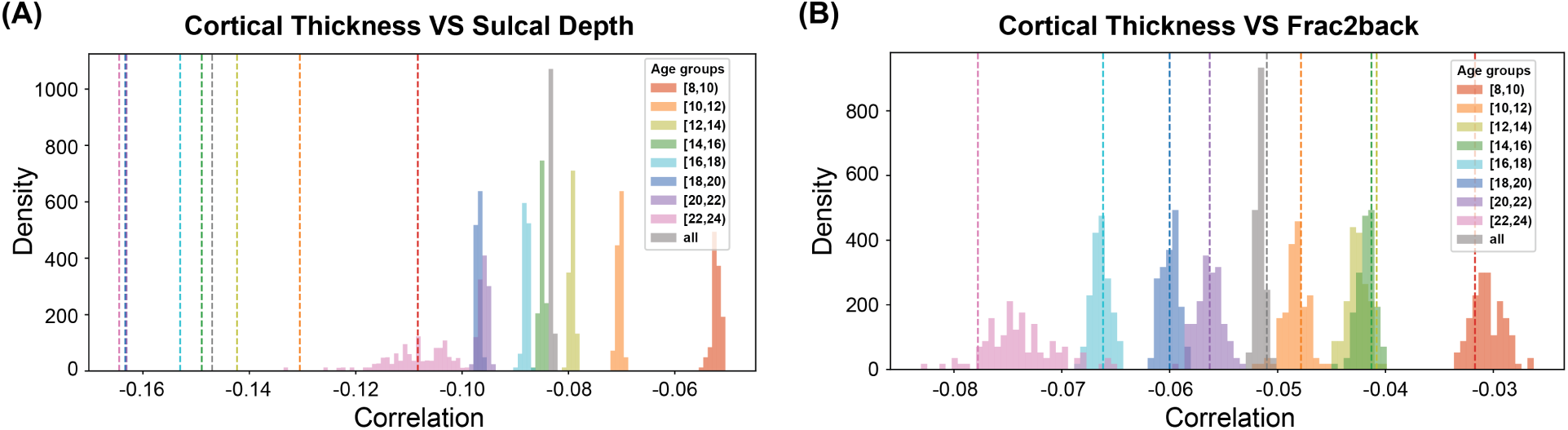
Null distributions (histograms) and the observed values (dashed lines) of the correlation test statistic obtained from permutations in global and conditional SPICE tests on the PNC dataset.

When performing conditional SPICE tests in each age group, all of the results agreed with the global results (Table 1) – the null hypothesis was rejected for cortical thickness and sulcal depth but not between cortical thickness and Frac2back. While in this population, the subject-level effect is strong enough that similar results are reached through global and conditional SPICE, these analyses revealed a potentially confounding interaction between the subject-level covariate age and the intermodal correspondence of interest. Namely, age systematically alters the marginal distribution of one or both modalities, such that the rejection reflects a mixture of genuine within-subject coupling and age-driven distributional shifts, rather than a homogeneous coupling effect across the population. The following example demonstrates how such confusion may arise in real data.

### 3.4 Global SPICE test produces conflicting results due to age effects

To further explore whether the coupling effects are homogeneous across the whole brain, we next performed the SPICE test on each of the 17 functional networks defined by Yeo et al. (2011). Specifically, we were interested in if intermodal correspondence exists at the parcel level, and how these measures may be influenced by development. As in the whole-brain analyses above, we compared the same two pairs of modalities. In addition, we considered pairwise combinations of age groups and applied the SPICE test to the combined sample in which subjects fell into either age range.

As speculated, we identified instances of Simpson’s paradox. Figure 2 provides an illustration using the comparison between cortical thickness and Frac2back within the Somatomotor B network. Without considering the age covariate, the SPICE test returned a spuriously small *p*-value on the combined sampling consisting of subjects aged 16 to 20, despite no subject-level correspondence being found within either age group ([16, 18) : *p* = 0.133; [18, 20) : *p* = 0.152; combined : *p* = 0.038). This disagreement persisted after accounting for sample size differences, where we resampled half of subjects from the mixed sample 1000 times and found the global SPICE test to reject the null in 28.2% resampled datasets.

**Figure 2:**
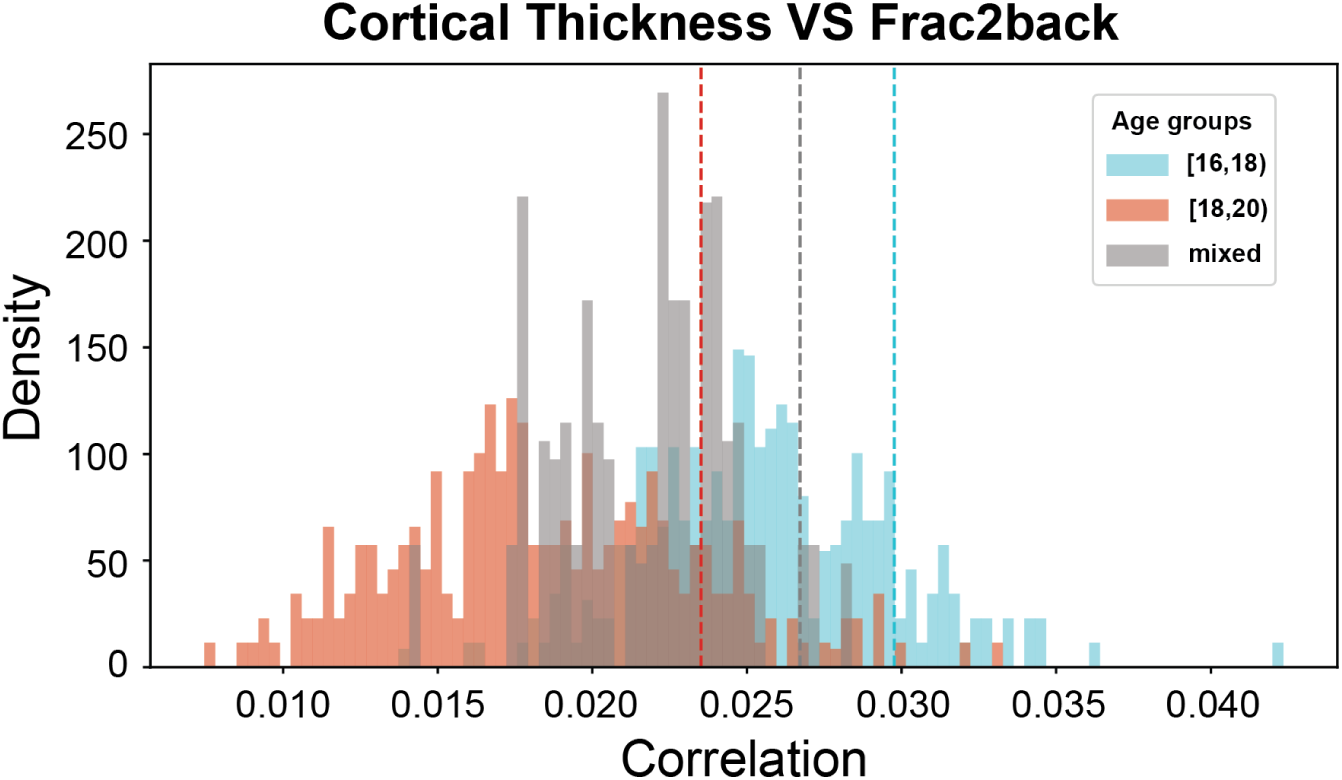
Confounding effect illustrated by the Somatomotor B network ([16, 18) : *p* = 0.133, [18, 20) : *p* = 0.152, mixed : *p* = 0.038∗).

Due to the unknown intermodal relationship between cross-group discordant pairs, there is no closed-form solution for the combined null distribution unless stronger homogeneity assumptions are made. In the following sections, we develop a hypothesis testing framework to detect such confounding covariates to avoid misinterpretation of SPICE test results.

## 4 Testing For Covariate Effects In The SPICE Test

### 4.1 Null hypothesis

We present a systematic approach to identify if covariate effects require conditional SPICE tests to be performed within subpopulations. If

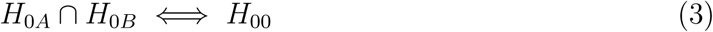

holds, then

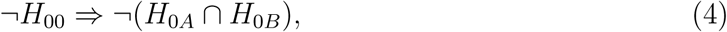

in which case a single SPICE test run with all subjects’ data would suffice to claim correspondence both at the population level and in at least one group. Thus, equivalence (3) eliminates ambiguity in the interpretation of test results due to confounding covariates. In reality, (3) does not necessarily hold. *H*_0*A*_*, H*_0*B*_ do not involve any statement about the distribution of *ψ*(*X_i_, Y_j_*) ∀*i* ∈ *A, j* ∈ *B*, which constitute a necessary portion of the right hand side of *H*_00_. This gap is naturally revealed by the SPICE procedure specifically due to its permutation-based estimation of the null distribution – shuffling within each group separately versus across all subjects freely.

Therefore, to bridge the global and conditional SPICE hypotheses, we need to assess an additional condition on the distributions of the between-group versus within-group intermodal correspondences. Formally, we define the difference-by-group null hypothesis as

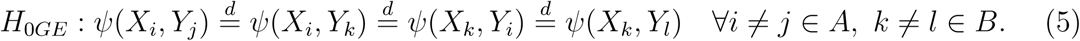

This strong null hypothesis states that the distribution of intermodal correspondence between two maps from different individuals does not depend on their covariate value (which group they belong to). Testing for *H*_0_*_GE_* allows us to detect if the impact of a covariate is sufficiently strong to alter the interpretation of group-level coupling. Theorem 4.1 describes the relationship between the global null, *H*_00_, and the conditional nulls, *H*_0_*_A_, H*_0_*_B_*. See Appendix for the proof.

#### Theorem 4.1

*If H*_0*GE*_ *holds, then*

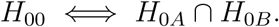

*Therefore, if intermodal correspondence exists at the global scale, then it must also exist in at least one of the sub-populations*.

*If H*_0_*_GE_ does not hold, then*

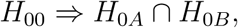

*But*

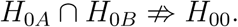

*Therefore, the two modalities may not exhibit any correspondence even in the presence of a marginal population-level relationship*.

A valid hypothesis testing procedure for *H*_0_*_GE_* would guide an analyst on whether the SPICE test should be performed in a covariate-aware manner. If *H*_0_*_GE_* holds, then results from the global SPICE test provide information about both group-specific and population-level relationships. If *H*_0_*_GE_* does not hold, then measurements from the two groups are sufficiently different that group assignment may drive spurious correlations between the modalities. The SPICE testing should be conducted within groups to account for such covariate effects (Figure 3).

**Figure 3:**
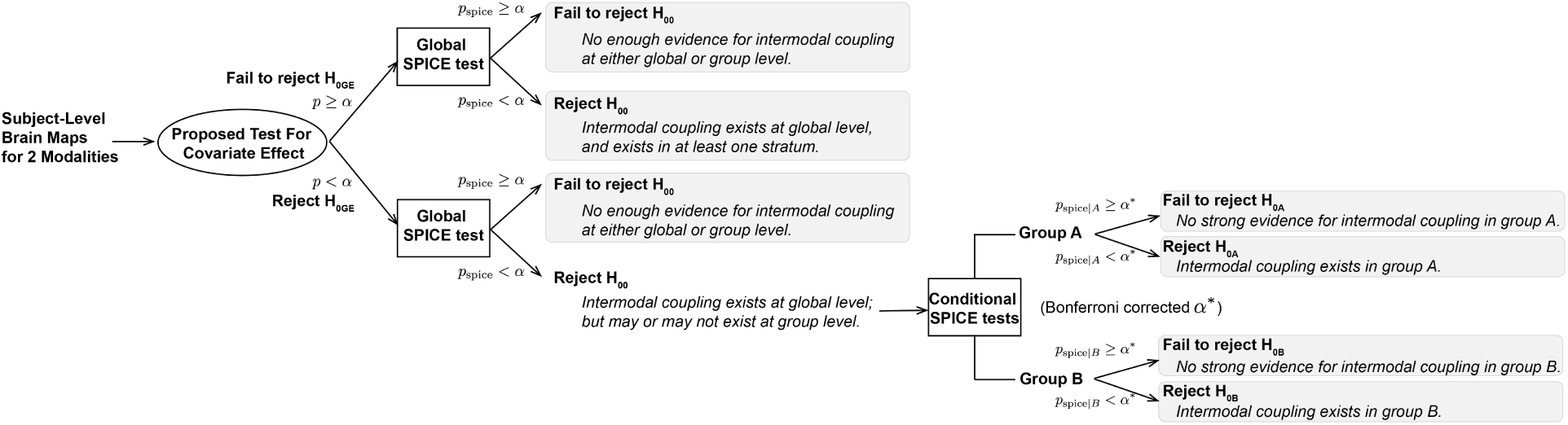
A covariate effect-aware workflow for performing the SPICE test.

### 4.2 Test statistic

Intuitively, a desirable test statistic for the difference-by-group null hypothesis *H*_0_*_GE_* should highlight potential differences between the within-group versus cross-group discordant correspondences. First, to simplify the notation, we define the set of concatenated map vectors for each individual to be

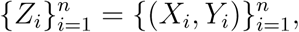

and define a symmetric kernel:

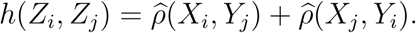

We define a test statistic for *H*_0*GE*_ in the form of a difference between U statistics:

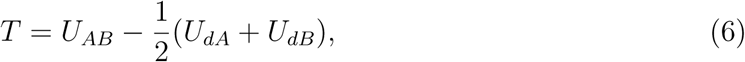

where

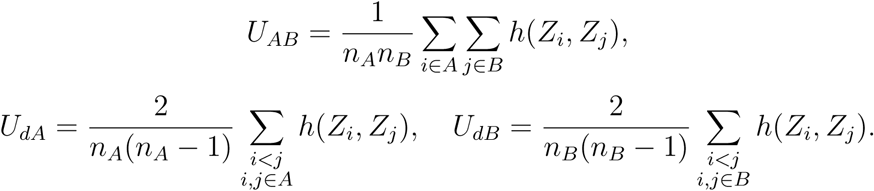

By definition, *h*(*Z_i_, Z_j_*) measures the strength of intermodal correspondence between maps from different individuals. Thus, *U_AB_* estimates this discordant correlation when the two individuals come from different groups; whereas *U_dA_* and *U_dB_* average this discordant correspondence within each group, respectively.

Intuitively, if the brain structure differs quite apparently between two groups of people, we expect that intermodal correspondence between the brain maps of two individuals randomly selected from within a group would be stronger than the cross-group correspondence. For example, one would expect males to be most similar to males (and females to females), whereas the male-to-female correspondence would be lower in magnitude. As defined in (6), the test statistic *T* is sensitive to such deviations in the average intermodal correspondence for between-group versus within-group discordant pairs.

### 4.3 Asymptotic distribution

Theorem 4.2 states the asymptotic distribution of the test statistic T under the null hypothesis *H*_0_*_GE_*. The full proof is given in the Appendix.

#### Theorem 4.2

*Define the integral operator*

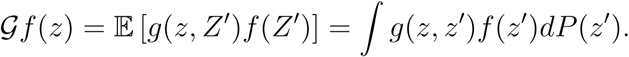

*Let η*_1_*, η*_2_*, … be the eigenvalues of* G*, with orthonormal eigenfunctions ϕ*_1_*, ϕ*_2_*, …:*

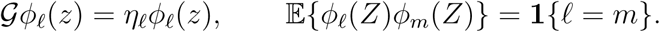

*Corresponding to each eigenvalue η_ℓ_, let* 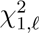 *be a chi-square random variable. Let* 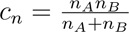.

*Assume that as* 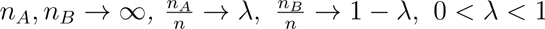*. Then,*

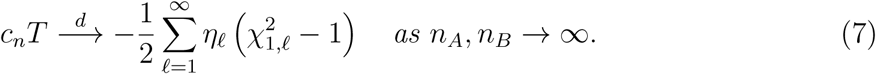

### 4.4 Hypothesis testing

The asymptotic distribution of the test statistic *T*, given in (7), is a random sum that involves an infinite number of eigenvalues. For estimation and hypothesis testing, we construct the *n* × *n* empirical kernel matrix *H* = {*H_ij_*}_1≤*i*≠j≤*n*_ where

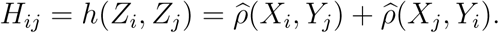

This empirical kernel matrix can be centered by 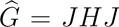, where 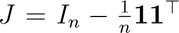. Let 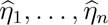 be the eigenvalues of 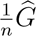. Then the limiting null distribution of *c_n_T* can be estimated by simulating *B* times

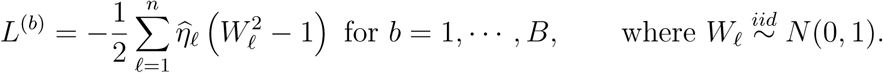

To test for *H*_0*GE*_, we compare the observed statistic *c_n_T* against the empirical CDF obtained from *L*^(*b*)^, with 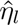 estimated from the data. For a one-sided test, the *p*-value is given by

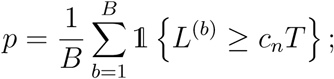

and for a two-sided test,

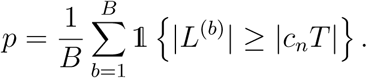

## 5 Simulations

### 5.1 Fully synthetic simulations

#### 5.1.1 Data generation

To assess the validity of the proposed test for covariate effects in the SPICE test, we evaluated its performance with data simulated in a controlled setting. In particular, we included a covariate that confounded intermodal coupling to varying degrees, in addition to a homogeneous nuisance covariate.

Let *i* index individuals for which *p*-dimensional brain maps are generated. Let *g_i_* ∈ {*A, B*} be a confounding covariate in the intermodal correspondence analysis, and *h_i_* ∈ {*a, b*} be a nuisance variable. Let *n* be the total sample size, *n_g,h_*be the number of individuals wit covariates *g_i_, h_i_*, and 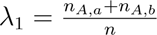 and 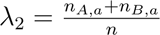 be the marginal fractions for the two variables, respectively. Consider the following data generating process:

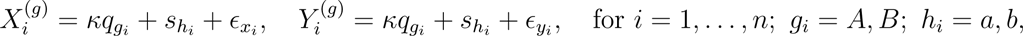

where 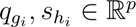 are standardized vectors with mean of 0 and variance of 1, and 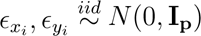 introduce individual-level randomness. To create group effects, let corr(*q_A_, q_B_*) = *r <* 1. The second covariate is nuisance, defined by *s_a_* = *s_b_* = *s*. Then, given fixed *n, p, λ*_1_*, λ*_2_, the group effect due to the first covariate *q_g_* should increase as *r* decreased from 1 (no group effect) to 0, with a signal-to-noise ratio controlled by *κ*. No group effect was expected with respect to the second covariate 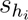.

We swept over a grid of parameter values (Table 2) to evaluate the Type I error rate for the nuisance covariate 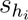 and the statistical power for 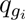, where rejection rates were calculated over 1000 simulations.

**Table 2:** Parameter values used in simulations.

| Parameter | Values |
| --- | --- |
| $r$ | 0, 0.2, 0.4, 0.6, 0.8, 1 |
| $\kappa$ | 0.2, 0.4, 0.6, 0.8 |
| $n$ | 100, 200, …, 1000 |
| $\lambda_1, \lambda_2$ | 0.25, 0.5, 0.75 |
| $p$ | 100, 1000 |

#### 5.1.2 Results

Regardless of the level of confounding effects in the first covariate 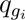, Type-I error rates were preserved around or below the nominal rate of 0.05 for *H*_0_*_GE_* regarding the nuisance variable (Supplementary Figure 1), indicating desirable performance. For the first covariate with a confounding effect, we evaluated power using cases with *r <* 1, where maps from the two groups were not perfectly correlated. A smaller *r* leads to groups that are more distinguishable. The rejection rates stayed at 1 for all but one setting where sample size was small (*n* = 100: rejection rate = 0.95; Supplementary Figure 2).

### 5.2 Plasmode simulations

#### 5.2.1 Data generation

Through the parametric simulation studies, we demonstrated that the proposed test can effectively detect covariate effects in the SPICE test with appropriate Type-I error control and sufficient power. Next, we conducted a plasmode simulation study to assess the performance of the test when applied to data that possess similar structure to real-world neuroimaging data.

We created semi-synthetic neuroimaging data based on cortical thickness and sulcal depth maps from the PNC. Vandekar et al. (2016) showed that the local correspondence between cortical thickness and sulcal depth differs between males and females, with females exhibiting stronger negative cortical thickness–sulcal depth coupling in several association cortices. Motivated by this finding, we simulated data leveraging this sex effect. For each subject, we generated pseudo-cortical thickness and sulcal depth data by adding i.i.d. Gaussian random noise to the normalized average maps of the sex group to which the subject belongs:

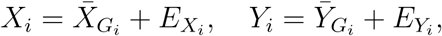

where 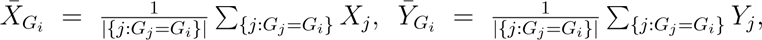 and 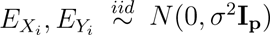. By construction, within-subject intermodal coupling was suppressed, while a strong sex effect was introduced.

We varied the noise level *σ*^2^ and ran 1000 simulations for each setting. In each simulation, we performed the test for covariate effects, which allowed us to select the appropriate SPICE tests to run. Additionally, we compared our approach to a naive global SPICE test, considered as a baseline approach.

#### 5.2.2 Results

The test correctly rejected the difference-by-group null *H*_0_*_GE_* in every simulation, suggesting the need to run conditional SPICE tests on each group separately. These conditional SPICE tests properly maintained Type I error rates as expected (omitted). Notably, the baseline approach of global SPICE tests showed liberal rejections especially when the data contained less noise, despite our intentional removal of subject-level coupling (Figure 4). Naive to confounding effects, this approach failed to distinguish the large-scale difference in groups from the true underlying subject-level coupling effect.

**Figure 4:**
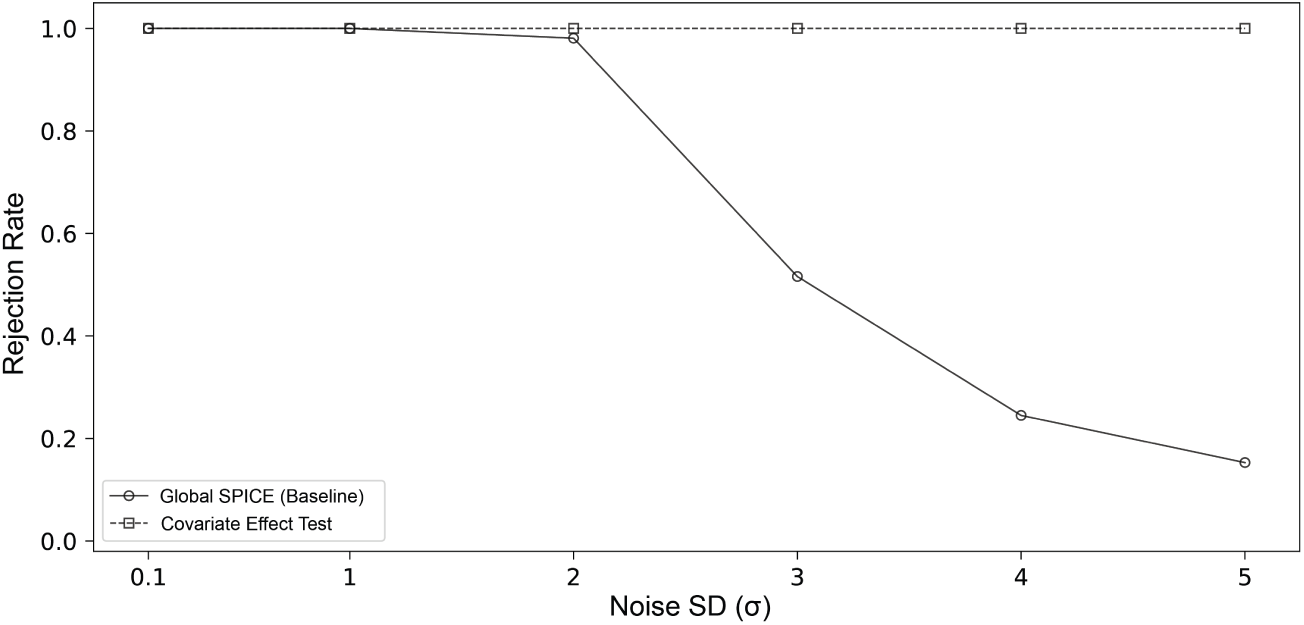
Rejection rates calculated from 1000 plasmode simulations under each parameter setting. The global SPICE test falsely rejected the SPICE null in the absence of true subject-level intermodal correspondence (round, dashed); our proposed test appropriately rejected the group effect null in every simulation across different noise levels (square, solid).

## 6 Application to the PNC Dataset

We now return to the PNC dataset (as in Section 3); in addition to the previously described age groups, we also performed a stratification by sex (617 males, 730 females). To highlight the utility of the proposed methods, we analyzed example subpopulations.

First, we considered the comparison of two age groups, [8, 10) and [18, 20). Our proposed test rejected the difference-by-group null hypothesis *H*_0*GE*_ for both comparisons between cortical thickness and sulcal depth (*p* = 0.00026∗) and between cortical thickness and Frac2back (*p* = 0.00012∗). Thus, we performed conditional SPICE tests within each age group separately (as reported in Table 1).

Next, we examined the binary covariate, sex. For both pairs of modalities, our test for covariate effects rejected the null *H*_0*GE*_ (*p* = 0.00002∗), suggesting that brain maps from male and female participants may be sufficiently different to drive a population-level intermodal coupling. This result is consistent with existing literature documenting sex differences in brain morphology. Males and females differ systematically in cortical thickness and sulcal/gyral folding patterns, with females tending to show relatively thicker cortex in some regions after adjusting for total brain size, and males and females differing in sulcal depth and gyrification indices that emerge over adolescent development (Luders et al., 2006; Im et al., 2006; Ritchie et al., 2018). Because cortical thickness and sulcal depth are both shaped by these sex-differentiated maturational processes, sex could plausibly act as a shared driver of between-group differences in the marginal distributions of both modalities, thereby inflating the apparent population-level coupling even if within-subject coupling is similar across sexes.

We performed conditional SPICE tests for each of the two sex groups. For both the female and male groups, we found significant correspondence between cortical thickness and sulcal depth (*p <* 0.001∗), whereas we failed to reject the conditional null hypotheses for coupling between cortical thickness and Frac2back (*p* = 0.620, 0.948) as expected. These results were consistent with the global SPICE test. Although sex modifies the marginal distributions of cortical thickness and sulcal depth, it does not, in this particular example, induce a spurious within-subject coupling between structural and functional modalities beyond what is captured by the global test.

## 7 Discussion

The SPICE test is a powerful tool to quantitatively evaluate whether a pair of brain maps correspond to each other in a meaningful way. However, like many statistical tests, it suffers from Simpson’s paradox when confounding variables are not addressed. We demonstrated how analyses using the SPICE test naive to covariate effects may lead to improper interpretation of intermodal correspondence. Leveraging the properties of U-statistics, we proposed a formal hypothesis testing procedure to test whether a binary covariate induces significant differences in intermodal correspondence. Beyond its role as a safeguard against spurious inference, this test also introduces opportunities for scientific discovery of biologically meaningful moderators of intermodal coupling effects.

In both simulations, plasmode simulations, and real data, our proposed method achieved desirable Type I error control and statistical power across different data settings. With the real data, we re-evaluated the hypotheses tested in the paper which originally formulated SPICE (Weinstein et al., 2021). In this context, the subject-level correspondence between cortical thickness and sulcal depth was strong enough that the null hypothesis was rejected no matter which testing procedure was performed. Nonetheless, we were able to find cortical regions where spurious results were generated by the global SPICE test. More generally, we expect that other situations where the subject-level effect is more marginal will particularly benefit from consideration of conditional SPICE. Overall, our results represent an important evaluation and extension of the SPICE test that increases its rigor and generalizability.

Several practical implications follow from these results. In homogeneous samples, or when the scientific target is explicitly a marginal population-level correspondence, the original SPICE test remains valid and interpretable. In heterogeneous samples, however, investigators should examine whether key biological or technical covariates alter the level of intermodal correspondence before drawing mechanistic conclusions. Reporting both overall and subgroup-specific correspondence estimates may help distinguish broad population-level effects from covariate-specific patterns. In fact, beyond a statistical nuisance to be controlled, such a covariate may reveal biological mechanisms by which demographic or clinical factors shape the functional or structural architecture of the brain.

This work also suggests several directions for future research. First, the current procedure focuses on a binary covariate, which provides a useful starting point but does not capture the full complexity of covariate structure in modern neuroimaging studies. Although we only included a pair of age groups in Section 6, age is intrinsically continuous and the current framework does not accommodate such a covariate. Many important sources of heterogeneity, such as head motion, symptom severity, disease status, and scanner- or site-related measurements, are also continuous or multi-level and can systematically influence neuroimaging phenotypes (Van Dijk et al., 2012; Power et al., 2012; Pomponio et al., 2020; Fortin et al., 2018). Large-scale multimodal datasets also contain multiple potentially interacting covariates. Extending the proposed U-statistic framework to continuous and multiple covariates would therefore broaden its applicability. Prior work on adjusted U-statistics shows that reweighting strategies based on propensity or stratification scores can account for confounding covariates in nonparametric multi-sample testing (Satten et al., 2018), suggesting one possible direction for generalizing our approach beyond the binary setting.

Second, future work could move beyond detecting covariate-induced confounding toward developing a covariate-adjusted SPICE test that estimates residual intermodal correspondence after removing or balancing observed covariate effects. This problem is not solved by directly importing standard residual-permutation methods from the general linear model, such as the Freedman–Lane procedure, because those methods are designed for testing regression coefficients after accounting for nuisance regressors, whereas the SPICE test defines its null by breaking cross-modality subject pairings (Freedman and Lane, 1983; Winkler et al., 2014; Weinstein et al., 2021). A covariate-adjusted SPICE procedure would need to preserve the native spatial organization of each subject’s maps and the permutation logic, while ensuring that permutations compare subjects with comparable covariate profiles or operate on appropriately adjusted correspondence measures.

Third, a closer examination of our test results reveals an interesting aspect of intermodal coupling for future neuroscience studies to explore. When the coupling effect differs between two groups of people, one would expected the cross-group coupling to be weaker than within-group ones. However, we found in real data analysis that the cross-group correlations often lie between the two within-group correlations (e.g., Figure 2). This observation suggests that the brain map of one modality for a given subject might be better predicted by the brain map of the other modality from a person in the different group than that from someone in the same group. Future work may investigate if this phenomenon is specific to certain correlation metrics or if biological explanations can be provided. For example, one could posit that one modality at an early age predicts the map of a second modality at a later age (better than the within age-group). While this hypothesis is distinct from the subject-level correspondence for which SPICE is designed, this example highlights the new neuroscience questions that are enabled by testing procedures that explicitly consider group effects within a spatial correspondence framework.

In conclusion, we examined the effect of covariates on the SPICE test for intermodal correspondence, and found that Simpson’s Paradox can result in fictitious findings. We derived a testing procedure to identify such settings, and demonstrated that SPICE can be conducted after group stratification to yield unbiased results. We hope that these methods will add rigor to intermodal neuroscience research and enable new avenues for the study of neurodevelopment and disease.

## Acknowledgments

(R.S.) Research reported in this publication was supported by the National Institute of Mental Health, the National Institutes of Health, under award numbers R01MH112847 and R01MH123550. The content is solely the responsibility of the authors and does not necessarily represent the official views of the National Institutes of Health.

B.W. received funding from the Research Institute of the Children’s Hospital of Philadelphia.

## Data Availability Statement

Neuroimaging data from the Philadelphia Neurodevelopmental Cohort (PNC) is retrieved from the following URL: https://reprobrainchart.github.io/docs/get_data.

## Supplementary Materials

**supplementary_materials.pdf** Appendices (proof for Theorems presented); Supplementary Figures.

**Declaration of generative AI use** Claude (Anthropic, Sonnet 4.5) was used to improve the readability and language of the manuscript only. All scientific content, analysis, results, and conclusions are the authors’ original work. AI-suggested edits were carefully reviewed and revised, and the authors take full responsibility for the content of the published article.

