## supplementary_materials for "A Test for Confounding in Coupling of Multimodal Neuroimaging Data"

### Appendices

#### A Methods

##### A.1 Proof of Theorem 4.1

Recall that the global null in the SPICE test is

$$H_{00} : \psi(X_i, Y_i) \stackrel{d}{=} \psi(X_i, Y_j) \sim P_0 \quad \forall i \neq j.$$

The conditional null hypotheses for groups $A, B$ are

$$H_{0A} : \psi(X_i, Y_i) \stackrel{d}{=} \psi(X_i, Y_j) \sim P_{0A} \quad \forall i, j \in A;$$
$$H_{0B} : \psi(X_k, Y_k) \stackrel{d}{=} \psi(X_k, Y_l) \sim P_{0B} \quad \forall k, l \in B.$$

The difference-by-group null hypothesis is defined as

$$H_{0GE} : \psi(X_i, Y_j) \stackrel{d}{=} \psi(X_i, Y_k) \stackrel{d}{=} \psi(X_k, Y_i) \stackrel{d}{=} \psi(X_k, Y_l) \quad \forall i \neq j \in A,\ k \neq l \in B.$$

By definition,

$$H_{00} \Rightarrow H_{0A}, \quad H_{00} \Rightarrow H_{0B}$$

regardless of whether $H_{0GE}$ holds.

Now, to show under what condition the other direction $H_{0A} \cap H_{0B} \Rightarrow H_{00}$ holds, consider individuals $i \neq j \in A,\ k \neq l \in B$. Suppose $H_{0A}$ and $H_{0B}$ both hold. Without additional assumptions, maps may not be i.i.d. for individuals from different groups. Thus, $\psi(X_i, Y_i)$ and $\psi(X_j, Y_j)$ may not follow the same distribution, and neither do $\psi(X_i, Y_j)$ and $\psi(X_k, Y_l)$. $H_{00}$ does not necessarily hold.

To show that $H_{0A} \cap H_{0B} \cap H_{0GE} \Rightarrow H_{00}$, let $m \in A$ or $B$. Suppose that in addition to $H_{0A}, H_{0B}$, we have $\psi(X_i, Y_j) \stackrel{d}{=} \psi(X_i, Y_k)$. Then,

$$\psi(X_i, Y_i) \stackrel{d}{=} \psi(X_i, Y_m).$$

Similarly, let $\psi(X_k, Y_l) \stackrel{d}{=} \psi(X_k, Y_i)$ hold, then

$$\psi(X_k, Y_k) \stackrel{d}{=} \psi(X_k, Y_m).$$

Further, assume that $\psi(X_i, Y_k) \stackrel{d}{=} \psi(X_k, Y_i)$. Then,

$$\psi(X_i, Y_j) \stackrel{d}{=} \psi(X_k, Y_l),$$

yielding

$$\psi(X_i, Y_i) \stackrel{d}{=} \psi(X_i, Y_m) \stackrel{d}{=} \psi(X_k, Y_k) \stackrel{d}{=} \psi(X_k, Y_m),$$

which is equivalent to $H_{00}$.

#### A.2 Proof of Theorem 4.2

Under the strong null hypothesis $H_{0GE}$, all observed pairs of maps are i.i.d. across subjects:

$$Z_1, \dots, Z_n \overset{iid}{\sim} P.$$

Let $\theta = \mathbb{E}[h(Z, Z')]$, where $Z, Z' \overset{iid}{\sim} P$. Define the first-order projection

$$h_1(z) = \mathbb{E}[h(z, Z')] - \theta,$$

and the degenerate kernel

$$\begin{aligned} g(z, z') &= h(z, z') - \theta - h_1(z) - h_1(z') \\ &= h(z, z') - \mathbb{E}[h(z, Z')] - \mathbb{E}[h(Z, z')] + \theta. \end{aligned}$$

By construction, $\mathbb{E}[g(z, Z')] = 0$.

By the Hoeffding decomposition for order-two U-statistics (Hoeffding, 1948; Serfling, 1980), each U-statistic admits a first-order projection plus a degenerate second-order remainder. Specifically,

$$U_{AB} = \theta + \frac{1}{n_A} \sum_{i \in A} h_1(Z_i) + \frac{1}{n_B} \sum_{j \in B} h_1(Z_j) + U^g_{AB}, \quad \text{where } U^g_{AB} = \frac{1}{n_A n_B} \sum_{i \in A} \sum_{j \in B} g(Z_i, Z_j),$$

$$U_{dA} = \theta + \frac{2}{n_A} \sum_{i \in A} h_1(Z_i) + U^g_{dA}, \quad \text{where } U^g_{dA} = \frac{2}{n_A(n_A - 1)} \sum_{\substack{i<j \\ i,j \in A}} g(Z_i, Z_j),$$

$$U_{dB} = \theta + \frac{2}{n_B} \sum_{j \in B} h_1(Z_j) + U^g_{dB}, \quad \text{where } U^g_{dB} = \frac{2}{n_B(n_B - 1)} \sum_{\substack{i<j \\ i,j \in B}} g(Z_i, Z_j).$$

Then,

$$\begin{aligned} T &= U_{AB} - \frac{1}{2}(U_{dA} + U_{dB}) \\ &= U^g_{AB} - \frac{1}{2}\left(U^g_{dA} + U^g_{dB}\right). \end{aligned}$$

The constant terms and the first-order projection terms cancel. Thus, $T$ is a second-order degenerate U-statistic under $H_{0GE}$.

To analyze the asymptotic distribution with higher-order terms, define the integral operator

$$\mathcal{G}f(z) = \mathbb{E}[g(z, Z')f(Z')] = \int g(z, z')f(z')dP(z').$$

Let $\eta_1, \eta_2, \dots$ be the eigenvalues of $\mathcal{G}$, with orthonormal eigenfunctions $\phi_1, \phi_2, \dots$:

$$\mathcal{G}\phi_\ell(z) = \eta_\ell \phi_\ell(z), \qquad \mathbb{E}\{\phi_\ell(Z)\phi_m(Z)\} = \mathbf{1}\{\ell = m\}.$$

Then in $L^2(P \times P)$,

$$g(z, z') = \sum_{\ell=1}^{\infty} \eta_\ell \phi_\ell(z) \phi_\ell(z').$$

Let $c_n = \frac{n_A n_B}{n_A + n_B}$. Assume that as $n_A, n_B \to \infty$, $\frac{n_A}{n} \to \lambda$, $\frac{n_B}{n} \to 1 - \lambda$, $0 < \lambda < 1$. Then by Theorem 12.10 and Example 12.11 in van der Vaart (1998),

$$c_n T = \frac{n_A n_B}{n_A + n_B} T \xrightarrow{d} -\frac{1}{2} \sum_{\ell=1}^{\infty} \eta_\ell \left(\chi^2_{1,\ell} - 1\right) \quad \text{as } n_A, n_B \to \infty,$$

where $\chi^2_{1,\ell} = W_\ell^2, \qquad W_\ell \overset{iid}{\sim} N(0,1)$.

### Supplementary Figures

Results for simulations described in Section 5.1:

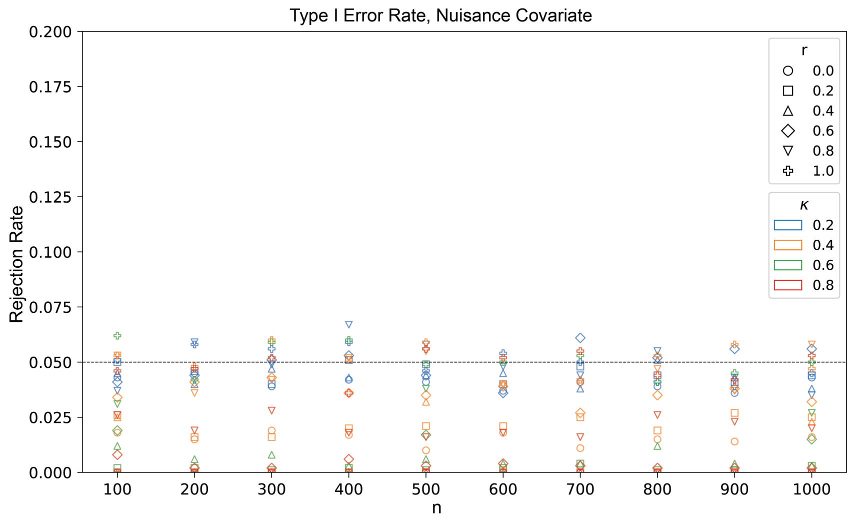


Figure 1: Type-I error rates for the nuisance covariate.

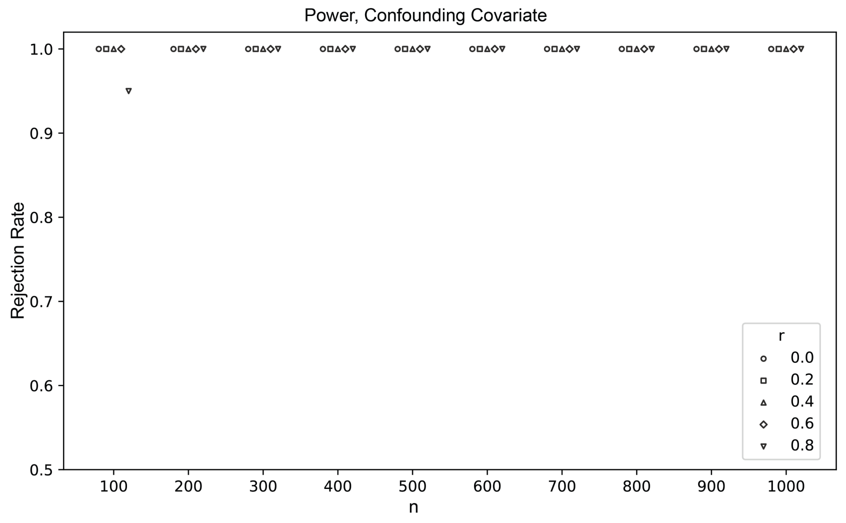


Figure 2: Power for the confounding covariate obtained under different effect levels (fixing $\kappa = 0.2, p = 100, \lambda_1 = \lambda_2 = 0.5$).
